# Comparative transcriptomics reveal contrasting fungal strategies in a plant pathogen versus an endophyte during initial host colonization

**DOI:** 10.1101/2024.12.17.628946

**Authors:** Soumya Moonjely, Frances Trail

## Abstract

Conidial germination marks the beginning of the fungal life cycle on the host plant, leading to disease or mutually beneficial relationships. Here, we use comparative transcriptomics to unravel the transcriptional similarities and differences during conidial germination and initial colony establishment of the plant pathogen *Fusarium graminearum,* and the endophyte *Metarhizium anisopliae*. Our comparison crosses four stages from fresh conidia to polar growth, hyphal extension, ending in either first hyphal branching (on medium) or appressorium formation (on barley). *F. graminearum* exhibited a higher number of upregulated genes for CAZymes, specialized metabolites and effectors compared to *M. anisopliae* during the interaction with the host, particularly during the appressorium stage, reflecting its pathogenic nature. The formation of appressoria by *M. anisopliae* conidia during germination on barley roots has not been documented previously and includes both morphological characteristics and gene expression patterns that regulate appressorium development. Our analysis reveals reduced transcript levels of CAZyme and specialized metabolite genes in *M. anisopliae* compared to *F. graminearum*, reflecting a less aggressive host penetration approach. The candidate genes associated with indole-3-acetic acid synthesis were upregulated in *M. anisopliae* during the appressorium stage, supporting its endophytic lifestyle, and suggesting that the fungus uses a phytohormone based strategy to interact with plant hosts. Collectively, our findings expand the transcriptome resources and provide valuable insights into the gene networks involved in conidial germination and initiation of infection in pathogenic versus endophytic fungi, as well as documenting appressorium formation for the first time, in the endophytic life cycle of *M. anisopliae*.

**IMPORTANCE:** Conidial germination is the initial step for fungal colonization in diverse environments. Here we examine the transcriptional similarities and differences in conidial germination and colony establishment of *Fusarium graminearum* and *Metarhizium anisopliae*, two fungal species belongs to the Order Hypocreales with distinct lifestyles. *F. graminearum* is a plant pathogen and the causal agent of Fusarium head blight on cereal crops, whereas *M. anisopliae* is an insect pathogen and root endophyte which forms beneficial associations with plants. We compared the transcriptome profiles of these species under two nutrient conditions across four stages of conidial germination. Our study shows that the expression profile of the genes encoding carbohydrate-active enzymes, specialized metabolites, and putative effectors varies between *F. graminearum* and *M. anisopliae*. The results of this study provide insights into gene networks associated with spore germination stages on the host in a pathogenic versus an endophytic fungus.

## INTRODUCTION

Conidial germination initiates the establishment of new fungal colonies. During germination (Figure 1), conidia break dormancy (Stage 1) through isotropic swelling to initiate metabolic activities (1–3); begin polar growth (Stage 2), characterized by the emergence of the germ tube; proceed through hyphal extension (Stage 3); and, depending on the substrate, extend hyphal growth through branching (Stage 4) or forming appressoria (Stage 4), specialized infection structures. Appressoria breach the physical barrier of the host epidermis, resulting in fungal entry into the inner host tissues (4). Several studies have reported the significant role of environmental factors, including nutrient availability, light, humidity and temperature, on conidial germination (5–8). Variation in the lifestyles of fungi also impact the process of conidial germination that culminates in appressorium formation (4, 9–11). Elucidating gene expression patterns involved in each stage is important for managing fungal growth on hosts.

**Fig. 1.**
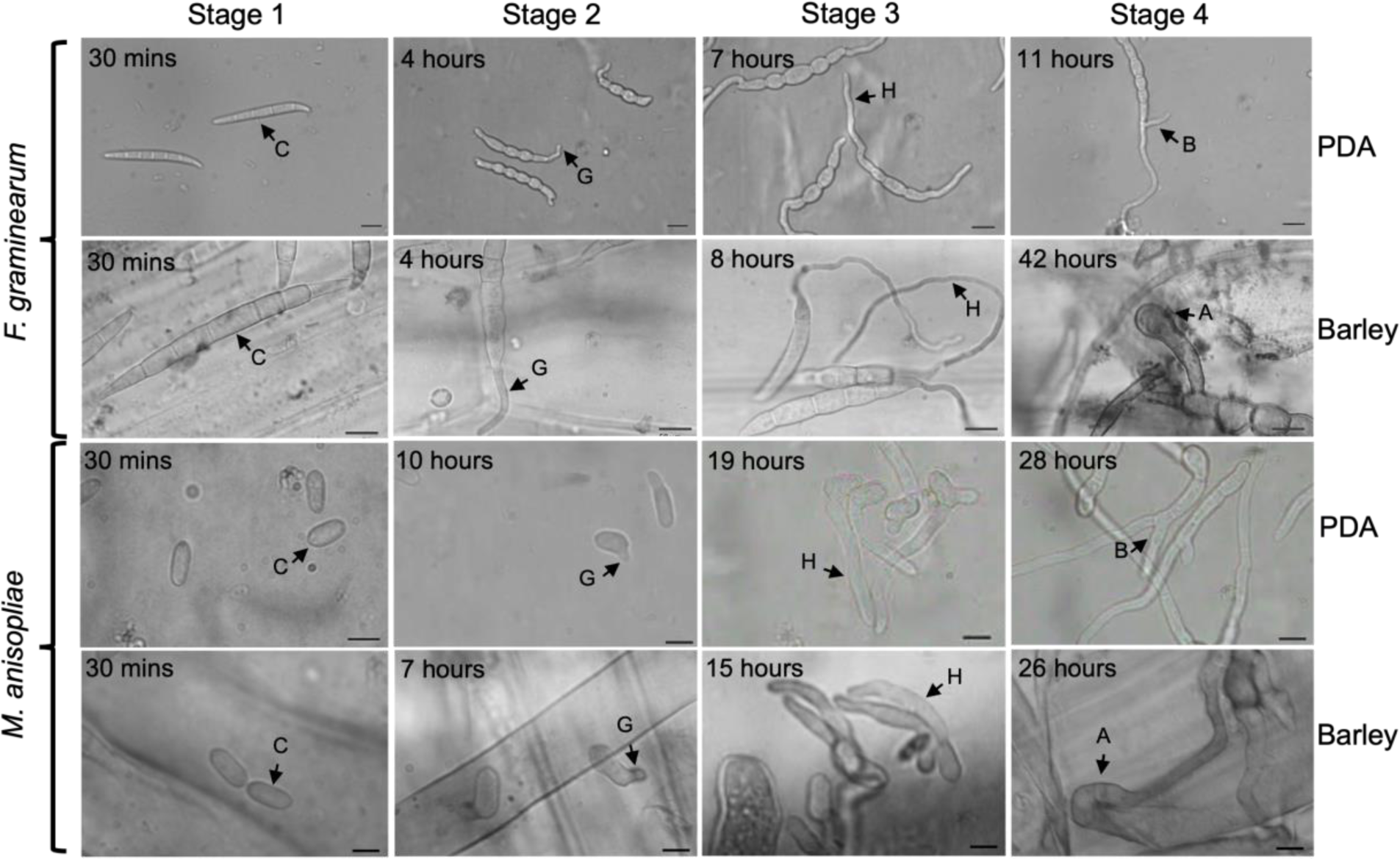
Time course and morphology of conidial germination stages in *F. graminearum* and *M. anisopliae* under two growth conditions on media and on barley. C – Conidium; G – Germ tube; H – Hypha; B-Hyphal branching, A-Appressorium. Scale bars: 20μm (*F. graminearum*); 5μm (*M. anisopliae*). Stage 1: Fresh conidia; Stage 2: Polar growth; Stage 3: Hyphal extension; Stage 4: First hyphal branching (PDA) or appressorium (barley).

Comparative transcriptomics is a valuable approach to gaining a deeper understanding of changes in gene expression patterns associated with alternate lifestyles among related fungal species. The technique provides information on the genetic basis of specific adaptations, such as those related to pathogenicity, symbiosis, and response to environmental stress. Additionally, comparative transcriptomics reveals how gene expression patterns have diverged or remained conserved across related species, providing insights into the molecular mechanisms of evolution and adaptation (12–15). Previously, we applied the technique to identify the molecular mechanisms involved in conidial germination *Fusarium graminearum* (Order Hypocreales) and *Magnaporthe oryzae* (Order *Magnaporthales*), both pathogenic to hosts in the Poaceae. The comparative transcriptomic approach revealed that the two species displayed similar gene expression patterns for the pre-penetration stages (16). However, substantial divergence in expression patterns between the two species was observed during appressorium formation. The study provided information on gene function associated with pathogenesis in fungal species of Orders *Hypocreale* and *Magnaporthales*. In the present study, *F. graminearum,* and *Metarhizium anisopliae* were compared, having different lifestyles, but both belonging to the Order *Hypocreales*.

Mutualistic and pathogenic fungi show distinct spore germination mechanisms on hosts, regulated by the activation of specific gene sets, due to their contrasting ecological roles and interactions with their plant hosts. Phytopathogenic fungi often express virulence factors and may employ strategies to weaken the host’s defense system, including producing enzymes, toxic metabolites, or invasive structures to breach the host’s barriers (17–22). Mutualistic fungi have evolved mechanisms to recognize and interact with their compatible hosts, often exhibiting a relatively passive colonization process where both partners benefit or only one partner benefits without harming the other (23, 24). Here, we analyze and compare the transcriptomic profiles of *F. graminearum* and *M. anisopliae* under two nutrient conditions (on medium and on the host) across four phases of conidial germination. *F. graminearum* is a mycotoxigenic phytopathogen that causes severe economic losses to cereal growers worldwide as the causal agent of Fusarium Head Blight (FHB) affecting wheat, barley, oats, and maize (25, 26). The FHB disease cycle is initiated when spores (conidia and/or ascospores) are deposited on the spikelets of a host plant, germinate and form infection structures (27). *M. anisopliae* is an endophytic fungus that lives inside plant tissue without causing disease symptoms (28, 29). In addition, *M. anisopliae* is a well-known insect pathogen with a wide host range (30, 31), and has the ability to transfer nitrogen from infected insects to plant hosts in exchange for carbon, thereby forming mutualistic associations with plant hosts (32, 33). While the process of conidial germination and subsequent penetration of insect hosts is well documented, there is little information on the gene expression changes during the initial stages of interaction with plants. Here we compare and contrast these two systems to better understand the specialization of each during spore germination on the host. For the host, barley leaf sheaths infected with *F. graminearum* germinating conidia and barley roots infected with *M. anisopliae* were studied, following the natural host of each fungus. We focused on the expression profile of the genes encoding carbohydrate active enzymes, specialized metabolites, and putative effectors across four stages of conidial germination, as these genes play critical roles in early fungal colonization on plants.

## MATERIALS AND METHODS

### Fungal isolates and experimental design

Wild-type (WT) *F. graminearum* (PH-1, NRRL31084) and *M. anisopliae* (ARSEF 549) cultures were routinely grown on V8 agar (Jackson and Bennett, 1990) at room temperature (RT, 22 - 24°C) and Potato Dextrose Agar (PDA, Neogen, Lansing, MI, USA) at 24°C, respectively. For both cultures, colonized agar blocks were stored long-term in 35% glycerol at -80°C. Conidia were generated for *F. graminearum* in carboxymethyl cellulose broth (CMC) in shaker culture in the dark for five days and then separated from hyphae by filtration through a double-layer of Miracloth (Sigma-Aldrich Inc, St. Louis, MO, USA). Conidia were collected by centrifugation, rinsed three times in sterile distilled water (dH_2_O), and resuspended in sterile dH_2_O to a final concentration of 1 x 10^6^ conidia/mL. For *M. anisopliae*, fresh conidia were produced in cultures grown in the dark on PDA for 10-12 days. Conidia were collected by flooding the plates with sterile 0.01% Triton X-100 in dH_2_O, rinsed, and adjusted to 1 x 10^6^ conidia/mL as for *F. graminearum*. For germination studies, PDA, which supports the conidial germination of both species, was used as a common garden environment. A 200 µL aliquot of the conidial suspension of each species was inoculated onto a sterile cellophane sheet (Research Products International, Mount Prospect, IL, USA) overlaid on PDA in a Petri dish (60 x 15 mm). Petri dishes containing inoculated sheets were then incubated at RT in continuous light (14-Watt, 6500 K, T12 white) till the conidia reached the appropriate germination stage. For *F. graminearum,* cellophane sheets from 30 plates were collected for each of three replicates, and total RNA was extracted from each germination stage. For *M. anisopliae,* cellophane sheets from 60 plates were pooled for RNA extraction for each replicate of each germination stage.

Barley leaf sheaths or roots were used to generate the transcriptome of conidial germination of the two species on the host. Leaf sheaths from 3-week-old barley seedlings were used to generate the transcriptomic profile of *F. graminearum* during initial pathogenesis. *Hordeum vulgare*, cultivar ‘Stander’, was grown in Suremix medium (Michigan Grower Products, Inc., Galesburg, MI, USA) under greenhouse conditions (22° C with a photoperiod of 16:8 h light:dark cycle). A conidial suspension of *F. graminearum* was prepared as described above, except the final concentration was adjusted to 2.5 x 10^6^ conidia/mL. To monitor germination stages, a 50 µL aliquot of conidial suspension was inoculated onto detached barley leaf sheaths (4-5 cm long) placed in a Petri dish lined with a moist filter paper to maintain humidity. The culture dishes were then incubated at RT in continuous light. Twenty barley leaf sheaths inoculated with *F. graminearum* were collected for each germination stage and used for total RNA extraction. Barley roots (cultivar ‘Stander’) were used to examine conidial developmental stages on the host. Surface sterilized barley seeds were germinated on 1% water agar. Germinated barley seedlings (4-5 days old) were transferred to a Petri dish, and 200 µL of the conidial suspension (2.5 x 10^6^ conidia/mL) was spread over the roots of each seedling and incubated at 24° C. The roots of five barley seedlings were collected for each stage and RNA was extracted. Each fungal germination stage on barley or PDA was examined and documented with light microscopy to confirm that at least 80% of the conidia reached the designated germination stage. Chlorazol black E and trypan blue were used to stain the *F. graminearum* and *M. anisopliae* cells on the host, respectively (16). Three independent biological replicates were prepared for each germination stage of *F. graminearum* and *M. anisopliae*.

### cDNA library preparation and RNA-sequencing

Total RNA of *F. graminearum* and *M. anisopliae* grown on cellophane membranes and on host tissue was isolated using Trizol reagent (Thermo Fisher Scientific, Waltham, MA, USA), as per manufacturer’s instructions. Harvested RNA was treated with RNase-Free DNase (Qiagen, Hilden, Germany), purified further using RNA Clean and Concentrator^TM^ - 5 (ZymoResearch, Irvine, CA, USA), and quantified using the Qubit™ Fluorometer (Invitrogen, Carlsbad, CA, USA). The integrity and quality of RNA samples were analyzed on High Sensitivity RNA ScreenTape (Agilent Technologies Inc., Santa Clara, CA, USA). The samples measuring with RNA integrity numbers greater than seven were submitted for RNA sequencing at the Research Technology Support Facility (RTSF) Genomics Core at Michigan State University (East Lansing, MI, USA, https://rtsf.natsci.msu.edu/). The cDNA libraries were prepared using the Illumina Stranded mRNA Library Prep Kit (Illumina, Inc., San Diego, CA, USA), and library sequencing was performed using either the HiSeq4000 (single-end, 50bp) or the NovaSeq6000 (single-end, 100 bp) Illumina (Illumina, Inc., San Diego, CA, USA) platform. Sequences were stored in FASTQ format.

### RNA-seq and differential expression analysis

The quality of the RNA-sequencing raw reads of transcriptome from conidial germination stages of *F. graminearum* and *M. anisopliae* were assessed with FastQC, and the low-quality reads were removed using the read filtering tool, Trimmomatic, Version 0.39 (34). The 3’ ends of NovaSeq6000 single-end 100 bp reads were trimmed to 50 bp reads (keeping the first 50 bp) using Trimmomatic Version 0.39. The reference genome assemblies of *F. graminearum* (35) and *M. anisopliae* (36) were obtained from EnsemblFungi (http://fungi.ensembl.org/). The trimmed reads were mapped to the indexed *F. graminearum* or *M. anisopliae* reference sequences using HISAT 2.2.1 (37). The HISAT alignments in SAM format were then sorted and converted to BAM format using SAM tools (38). Count files were then generated using HTSeq Version 0.11.1 (39). Gene expression patterns in germination Stages 2, 3 and 4 were compared to Stage 1. The DESeq2 package Version 1.30.1 in R was used to normalize the read counts and to perform differential expression analysis. Heatmaps were generated using pheatmap 1.0.12 and row z-scores were used to scale the expression between samples. For each species, the Differentially Expressed Genes (DEGs) for germination Stages 2, 3, and 4 compared to Stage 1 were identified using DESeq2. The genes with Log_2_ Fold Change (LFC) ≥ 2 and adjusted p-values (p-adj) ≤ 0.01 were designated as upregulated, and the genes with LFC ≤ -2 with p-adj ≤ 0.01 were designated as down-regulated. To further assess the variability among biological replicates, PCA was performed on the rlog-transformed values of transcriptome data of PDA and *in planta* samples.

### Annotation and databases

Carbohydrate-active enzymes (CAZymes) of *F. graminearum* and *M. anisopliae* were predicted using the dbCAN2 meta server (https://bcb.unl.edu/dbCAN2/index.php) HMMER: dbCAN, DIAMOND: CAZy and HMMER: dbCAN-sub packages (40). Proteins were predicted to be as CAZymes when at least two packages of the dbCAN meta server identified a query protein as a CAZyme, and these predicted CAZyme proteins were selected for further analysis. The dbCAN2 metaserver classified CAZymes according to their activity, such as glycoside hydrolases (GH; enzymes involved in the hydrolysis of glycosidic bonds), glycoside transferases (GT; enzymes involved in the formation of glycosidic bonds), polysaccharide lyases (PL, enzymes involved in the non-hydrolytic cleavage of glycosidic bonds), carbohydrate esterases (CE; enzymes involved in the hydrolysis of carbohydrate esters) and auxiliary activities (AA; redox enzymes that act in conjunction with CAZymes). In addition, a non-catalytic class of CAZymes, carbohydrate binding modules (CBM; adhesion to carbohydrates), has also been described (http://www.cazy.org). Specialized metabolite gene clusters were detected in *M. anisopliae* with the antiSMASH 6.0 (fungal version) web server (41). We examined the expression specialized (secondary) metabolite gene clusters that have been identified in the genome of *F. graminearum* previously (42–45). The selected CAZymes and specialized metabolite genes were compared with those identified in previous studies (36, 44) and other databases (http://www.cazy.org and https://mycocosm.jgi.doe.gov/mycocosm/home). EffectorP v 2.0 (46) was used to identify putative effector proteins. SignalP-5.0 was used to detect the presence of signal peptides. Transmembrane domains were detected using DeepTMHMM. We selected the genes with Log_2_ Fold Change (LFC) ≥ 5 and adjusted p-values (p-adj) ≤ 0.01 for CAZyme, specialized metabolite and effector gene expression analysis. Gene ontology (GO) (47) and Metacyc metabolic pathway analyses (48) were performed using FungiDB (49) with p-values <0.05 considered statistically significant.

### Data availability statement

The RNA-seq data generated in this work have been deposited in NCBI’s BioProject (https://www.ncbi.nlm.nih.gov/bioproject). The data is accessible through GEO series accession numbers <u>GSE277787</u> (conidial germination in *F. graminearum*) and <u>GSE277627</u> (conidial germination in *M. anisopliae*).

## RESULTS

### Conidial germination stages on PDA and the host for transcriptome analysis

Spore germination in *F. graminearum* and *M. anisopliae* Stages 1-4 were morphologically characterized and transcriptome profiles were completed on PDA and on barley (Fig. 1). A difference in developmental time-course between the growth conditions was not observed in Stages 2 and 3 in *F. graminearum.* At 11 hours (h) on PDA, the first hyphal branches were initiated, while the formation of infection structures on leaf sheaths was observed after 42h. The majority of *M. anisopliae* conidia reached Stages 2, 3 and 4 in PDA at 10h,19h and 28h, respectively, whereas on the host, the respective germination stages were reached by 7h, 15h and 26h.

### General analysis of the transcriptomes from conidial germination

The analysis of the transcriptomes of the conidial germination stages showed that the appressorium stages in *F. graminearum* and *M. anisopliae* were transcriptionally divergent compared to other germination stages. Principal Component Analysis (PCA) plots revealed that germination stages of *F. graminearum* (Fig. 2A) and *M. anisopliae* (Fig. 2B) were grouped separately on PDA and on barley, displaying the differences in transcriptome profiles between these artificial and natural conditions. Clustering also revealed similarity of gene expression profiles among biologically independent replicates, indicating a high level of replicability for the samples. The DEGs in *F. graminearum* and in *M. anisopliae* for all germination stages under both conditions are presented in Tables S1 and S2, respectively. The numbers of downregulated DEGs for all stages of *F. graminearum* grown on PDA were higher compared to those grown on barley. A higher number of upregulated DEGs compared to downregulated DEGs was observed specifically during the appressorium stage on the host (Figs. 2C, S1A-F) suggesting a dynamic gene activation process during host colonization. For *M. anisopliae* the number of upregulated DEGs was higher than the number of downregulated DEGs for all stages in both nutritional conditions (Fig. 2D, S2A-F). The numbers of shared and unique upregulated DEGs among the germination stages on PDA and barley (Fig. 3A-D) demonstrated that the appressorium stage is transcriptionally highly divergent from the rest of the germination stages in both species. We performed k-means clustering of DEGs to identify the top 100 genes that show variation in expression between two nutritional conditions. The appressorium stage showed a higher level of variation in the expression of genes in both species. In *F. graminearum,* the variable genes with higher expression levels were associated with specialized metabolite biosynthesis (Cluster 1) and carbohydrate metabolism (Cluster 3) (Fig. S3A). In *M. anisopliae*, cluster 4 contains the highly variable up-regulated genes for the appressorium stage that were associated with fatty acid metabolism, specialized metabolite biosynthesis, and effector protein encoding genes (Fig. S3B).

**Fig. 2.**
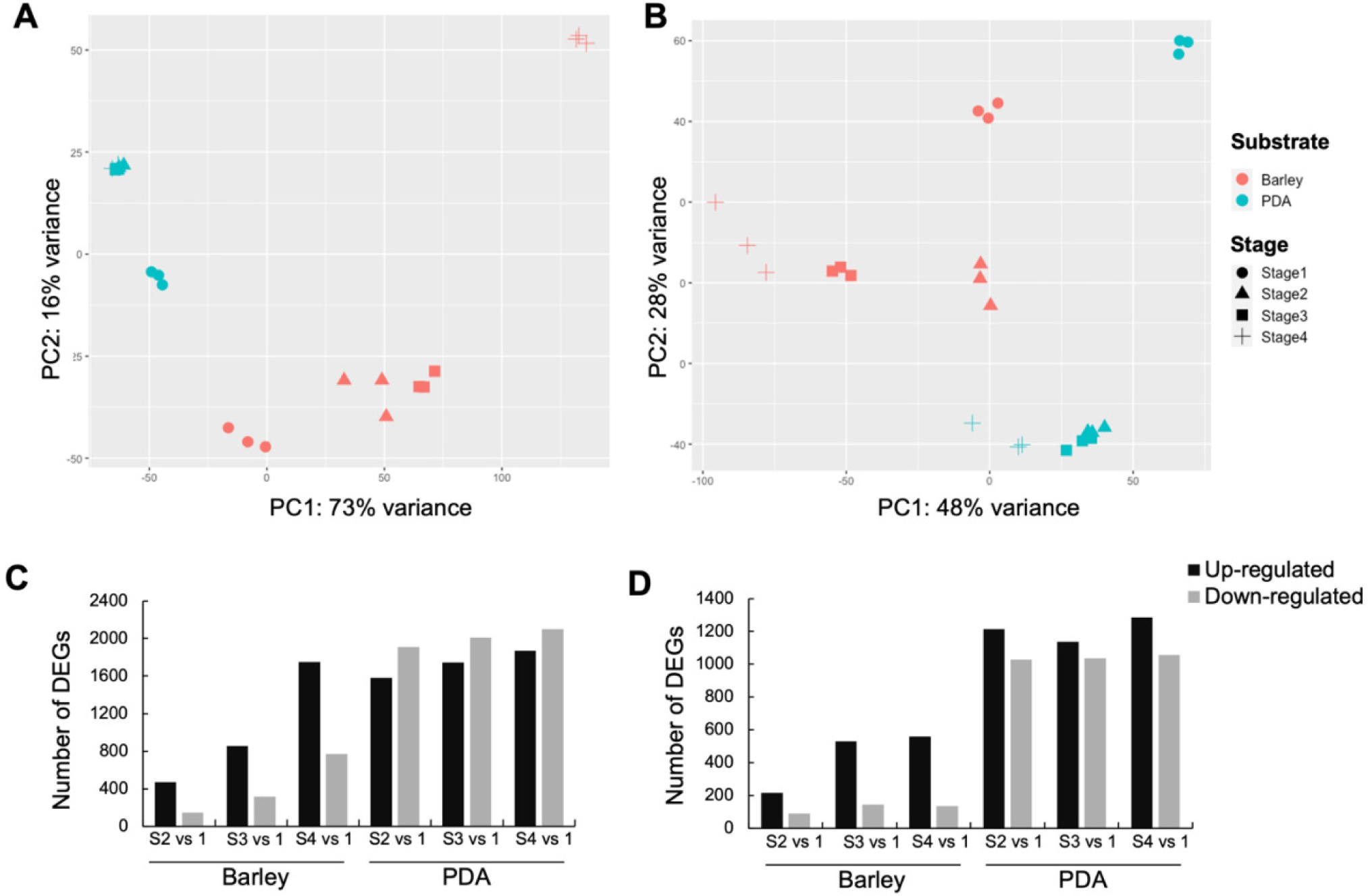
General analysis of the transcriptomes of conidial germination. The principal component analysis of the transcriptome data from germination stages of (**A**) *F. graminearum,* and (**B**) *M. anisopliae* on barley and PDA. The number of up- and down-regulated differentially expressed genes (DEGs) of (**C**) *F. graminearum* and (**D)** *M. anisopliae*.

**Fig. 3.**
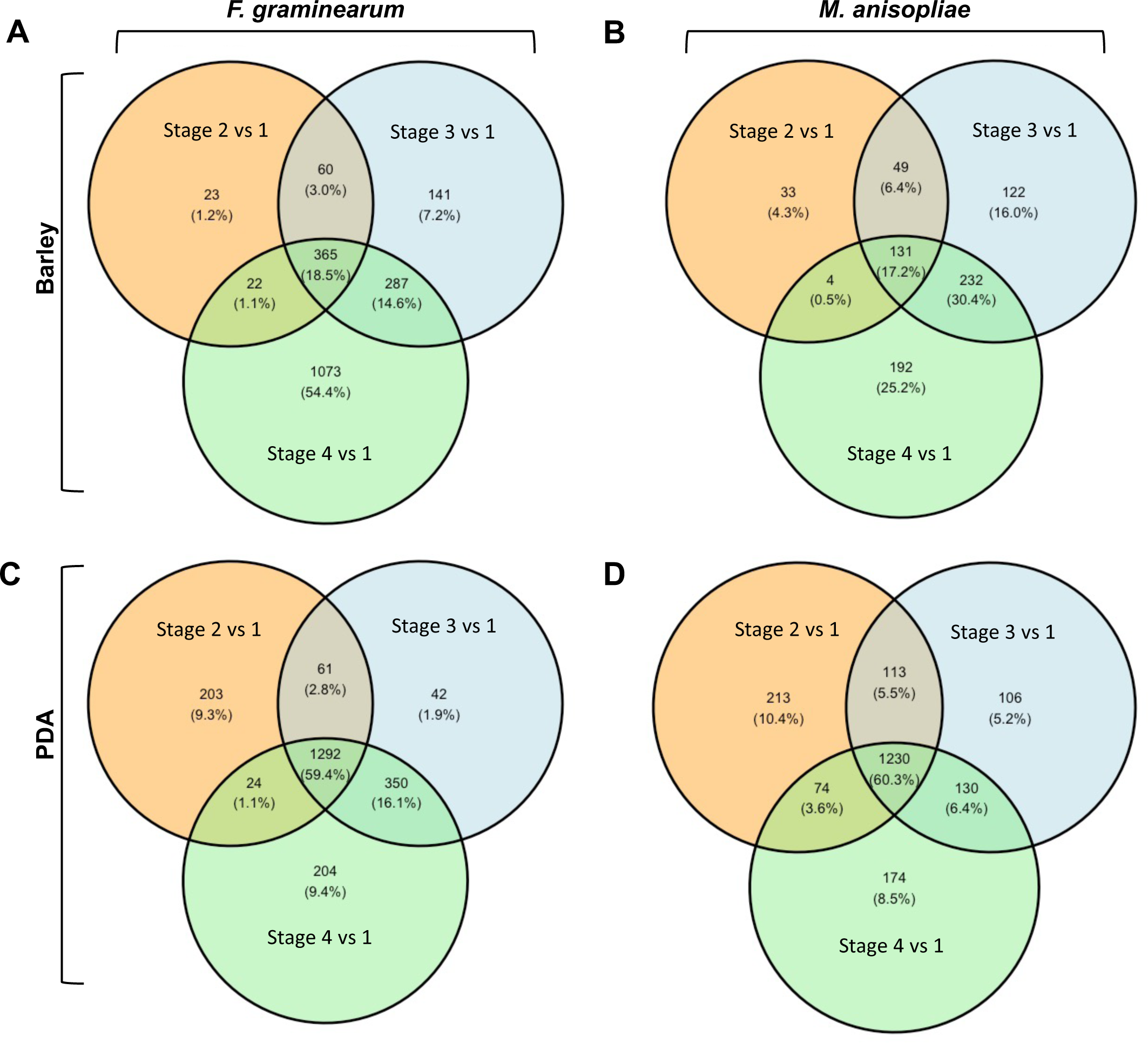
Shared and unique up-regulated DEGs (LFC ≥ 2 and adjusted p-adj ≤ 0.01) in the germination stages of *F. graminearum* and *M. anisopliae* on barley and PDA. (**A**) *F. graminearum* on barley; (**B**) *M. anisopliae* on barley; (**C**) *F. graminearum* on PDA and (**D**) *M. anisopliae* on PDA.

### Gene ontology and metabolic pathway analysis of upregulated DEGs

Gene Ontology (GO) and metabolic pathway analyses of *F. graminearum* (Fig. S4, Table S3 and S5) and *M. anisopliae* (Fig. S5, Table S4 and S6) revealed that during germination on the host, more carbohydrate breakdown and utilization genes were upregulated compared to expression of the same genes during germination on PDA, indicating a differential upregulation of CAZymes on the host in two fungal species. In *F. graminearum*, more carbohydrate breakdown and utilization genes were upregulated during conidial germination on the host, than in *M. anisopliae*, particularly those targeting plant cell walls. The enrichment of GO terms for ‘secondary metabolite biosynthesis’ during Stage 4 on the host supports a role for specialized metabolites during early stages of infection in *F. graminearum* (Fig. S4C). Genes related to appressorium development were also upregulated in *F. graminearum* at Stage 4 on the host. Notably, the ‘o-diquinones biosynthesis’ pathway was specifically enriched at this stage, including five tyrosinase-encoding genes (Table S5). Tyrosinases are components of the melanin biosynthesis pathway, where they contribute to both appressorium development and pathogenesis (50). GO enrichment analysis on PDA for both species showed the enrichment of GO term, ‘microtubule based movement’, suggesting the movement of organelles and microtubules was more active on the medium than they were during interactions with their natural host (Fig. S4D-F and S5D-F, Table S3 and S4). Interestingly, metabolic pathway analysis in *M. anisopliae* revealed that genes associated with IAA biosynthesis were upregulated during germination Stages 3 and 4 on barley. IAA biosynthesis genes were enriched in the L-tryptophan degradation pathways (PWY-6307 and TRYPDEG) (Table S6). Common metabolic pathways between the two species for each germination stage were examined (Fig. S6A-C, Table S7). During Stage 4 on the host, genes for degrading plant defense compounds were upregulated in both *F. graminearum* and *M. anisopliae*, demonstrating their adaptation for bypassing plant defenses during the appressoria formation (Table S7).

### Differential expression of CAZyme genes

GO enrichment analysis suggests that carbohydrate metabolism is highly active during germination stages of both fungal species on barley with evidence that the *F. graminearum* genome harbors 170 more genes encoding CAZymes than does *M. anisopliae* (Fig. 4A and B), and expression of CAZymes differed substantially between *F. graminearum* and *M. anisopliae* (Fig. 4C-F). *F. graminearum* expressed a higher number of upregulated CAZymes during germination on barley (13.4% (Stage 2), 21.9% (Stage 3), 31.4% (Stage 4) compared to PDA (4.6% (Stage 2), 4.8% (Stage 3), 5.7% (Stage 4) (Fig. 4C and D; Table S8), suggesting its ability to penetrate the host tissue and cause necrosis soon after the initiation of infection. In *M. anisopliae,* fewer CAZymes were differentially expressed overall (Fig. 4E and F; Table S9). During germination stages on the host, *M. anisopliae* expressed 2.5%, 5.6%, and 7.4% of CAZymes at Stages 2, 3, and 4, respectively, and the expression levels were 4.3%, 4.1%, and 4.9% at the same respective stages on PDA. However, a 1.5-fold increase in expression of glycoside hydrolases was observed during germination Stages 3 and 4 on the hosts compared to PDA. These results suggest a critical role for CAZymes in both pathogenic and endophytic species during early stages of colonization on the hosts. *F. graminearum* displayed upregulation of both chitin synthase and cutinase genes (Table S8), which was not observed in *M. anisopliae*. Among the PLs identified in *F. graminearum,* 80.3% were differentially upregulated during the appressorium stage on the host (Fig. 4G), suggesting the degradation of plant cell walls to initiate infection. The *M. anisopliae* genome harbors only two PL genes and one PL gene (MAN_03654) was found differentially upregulated in Stage 4 on the host. *F. graminearum* genome contains numerous genes for pectin-degrading enzymes which are differentially upregulated during germination on the host, especially in the appressorium stage. In *M. anisopliae*, the pectin-degrading enzyme genes were not upregulated during host colonization (Table S9), suggesting a minimal role of pectin degrading enzymes during the initial stages of host colonization.

**Fig. 4.**
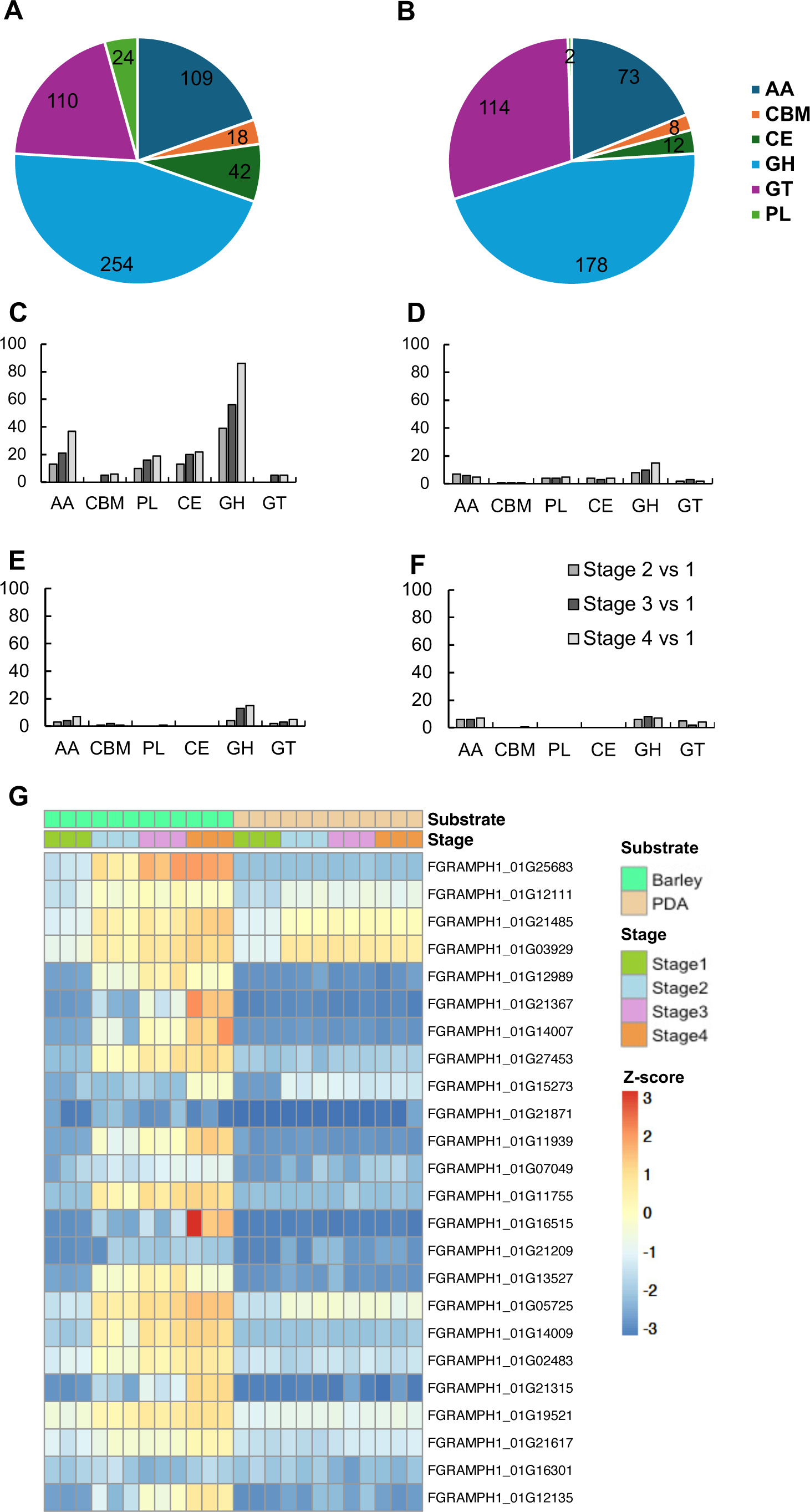
Differential expression of CAZyme genes in the germination stages of *F. graminearum* and *M. anisopliae*. CAZyme classes identified in the genome of (**A**) *F. graminearum* and (**B**) *M. anisopliae*. Number of differentially expressed genes encoding CAZymes during Stages 2, 3 and 4 in (**C**) *F. graminearum* on barley, (**D**) *F. graminearum* on PDA, (**E**) *M. anisopliae* on barley, **(F**) *M. anisopliae* on PDA. (**G**) Expression of polysaccharide lyase (PL) genes in *F. graminearum*. AA: Auxiliary Activities, CBM: Carbohydrate Binding Module, PL: Polysaccharide Lyases, CE: Carbohydrate Esterases, GH: Glycosyl Hydrolases, GT: Glycosyl transferases.

### Differential expression of specialized metabolite genes

We observed substantial differences in the expression of specialized metabolite genes between the two species during host colonization. During the appressorium stage in *F. graminearum*, the expression of specialized genes increased significantly, with a 3.4 to 6.8-fold upregulation compared to other germination stages (Table S10). The trichothecene biosynthetic gene cluster was upregulated during the appressorium stage on barley (Fig. S7, Table 1), likely producing trichothecenes in early host colonization events. In *M. anisopliae*, the 58.06% of specialized metabolite genes upregulated during the appressorium stage on the host were the same as upregulated on PDA (Table S11). Interestingly, the metabolic pathway analysis showed that genes associated with tryptophan metabolism were upregulated in Stages 3 and 4 on the host. Tryptophan is a precursor for the biosynthesis of phytohormone, IAA, that regulates various aspects of plant growth and development (51–53). *M. anisopliae* synthesizes indole-3-acetic acid (IAA), through tryptophan dependent pathways (54). Further analysis revealed that the candidate genes encoding enzymes associated with IAA biosynthesis, including anthranilate phosphoribosyltransferase, nitrilase and flavin monooxygenase, were upregulated in *M. anisopliae* in Stage 4 on the host (Table S12), supporting a role of IAA metabolism in the interaction between *M. anisopliae* and the host.

**Table 1.** The major specialized metabolite gene clusters upregulated in *F. graminearum* during germination stages on the host and PDA.

|  | Barley |  |  | PDA |  |  |
| --- | --- | --- | --- | --- | --- | --- |
|  | Stage<br>2 vs 1 | Stage<br>3 vs 1 | Stage<br>4 vs 1 | Stage<br>2 vs 1 | Stage<br>3 vs 1 | Stage<br>4 vs 1 |
| Orcinol/orsellinic acid | 2 | 3 | 4 | 4 | 4 | 4 |
| Triacetylfusarinine | 0 | 0 | 2 | 0 | 0 | 0 |
| Trichothecene | 1 | 2 | 11 | 1 | 2 | 1 |
| Butenolide | 0 | 1 | 1 | 0 | 0 | 0 |
| Precursor of insoluble perithecial pigment | 1 | 1 | 1 | 0 | 0 | 0 |
| Malonichrome | 0 | 0 | 3 | 0 | 0 | 0 |
| Decalin-containing diterpenoid pyrones | 0 | 0 | 7 | 0 | 0 | 0 |

The specialized metabolite genes with $LFC \geq 5$ and $p\text{-adj} \leq 0.01$ were selected for the analysis and the table shows the number of genes upregulated in each biosynthetic gene cluster.

### Differential expression of effector genes

Our results indicate that genes encoding secreted putative effectors are upregulated during the appressorium stages in both species, with *F. graminearum* expressing more effectors overall (Tables S13 and S14). The putative effector genes, AA1-like domain-containing proteins, LysM domain-containing proteins, and the cell wall mannoprotein 1 domain proteins were upregulated in both species during appressorium formation, emphasizing the pivotal role for secreted effector proteins in the early stages of plant colonization for both species. *M. anisopliae* upregulated putative effectors included a calcium channel inhibitor (Killer toxin Kp4) and a proteinase inhibitor I1 (Kazal domain) upregulated during the appressorium stage, indicating roles in modulating host immune responses during pathogen germination on the host. Besides the predicted effector genes, we also examined the expression of hydrophobin genes, adhesin genes, genes encoding proteins containing necrosis-inducing protein domains (IPR008701), and genes encoding proteins with Egh 16-like virulence factor domains (IPR021476), all of which are pathogenicity related genes that have been reported in other plant-colonizing. Our analysis suggests that the upregulation of hydrophobins during germination stages on the host indicates its role as a virulence factor during initial colonization of *F. graminearum* (Fig 5A). Additionally, the genes encoding proteins with Egh16-like domains, associated with some virulence factors, were upregulated during the appressorium stage in *F. graminearum* – a key stage in the infection process. However, in *M. anisopliae*, these genes were upregulated during Stages 2 and 3 (Fig 5B), when the fungal conidia were beginning to germinate, but their expression decreased as the fungus progressed to the appressorium stage.

**Fig. 5.**
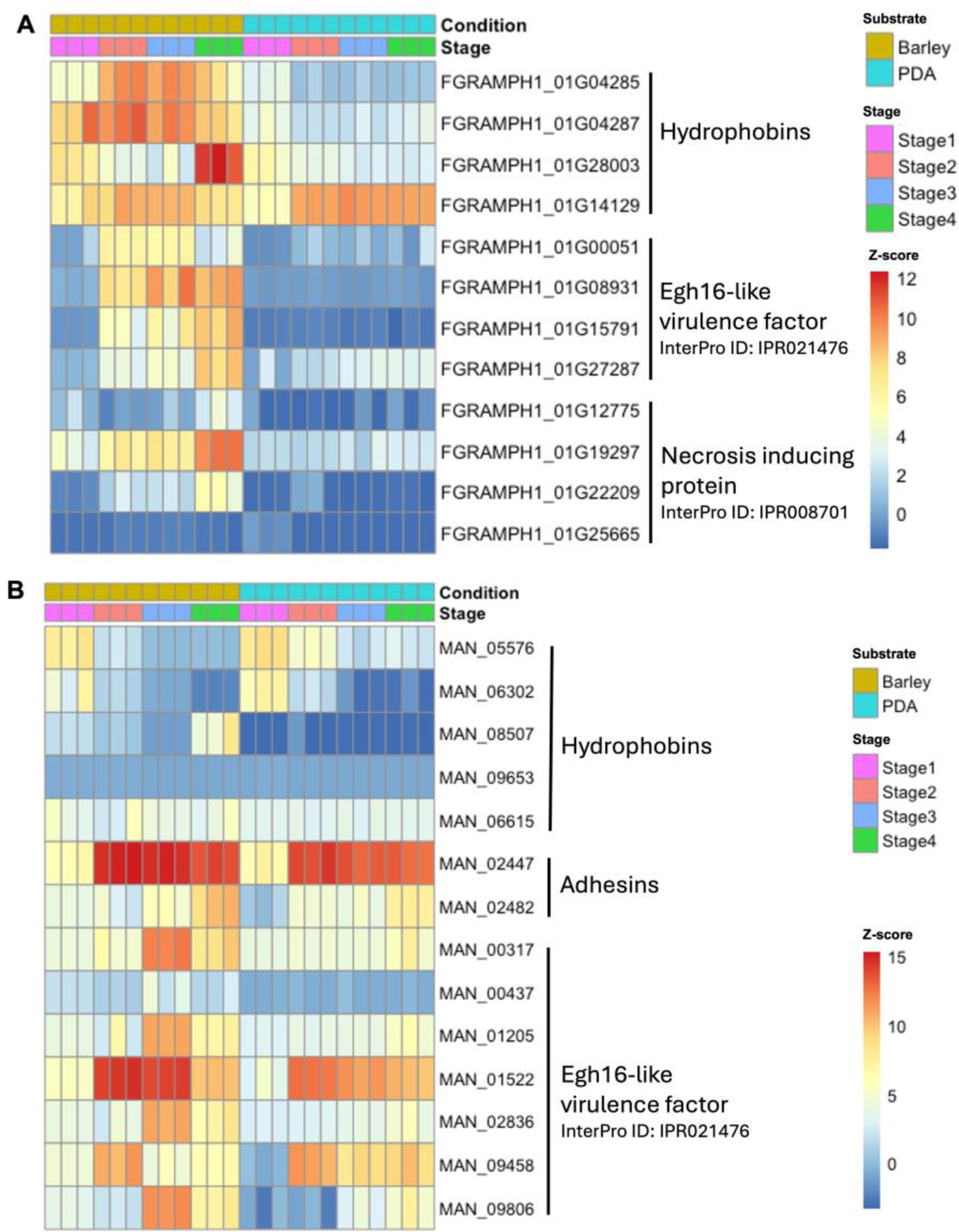
Expression of pathogenicity related genes in germination stages on barley and PDA. (**A**) *F. graminearum* and (**B**) *M. anisopliae.* Heatmap colour range represents high to low expression levels where the red represents higher expression and blue represents lower expression.

## DISCUSSION

We compared genome-wide expression of conidial germination stages of pathogenic versus. endophytic fungi on barley using *F. graminearum* and *M. anisopliae*. During the appressorium stage, both fungi exhibited a significant transcriptomic shift compared to other germination stages, highlighting the unique transcriptional changes associated with appressorium development. While *F. graminearum* exhibited a more aggressive colonization strategy, with extensive CAZyme and specialized metabolite involvement, *M. anisopliae* adopted a more subtle approach by expressing phytohormone biosynthetic genes and fewer CAZyme genes, to establish a symbiotic relationship with its host. Both fungi showed upregulation of the genes encoding LysM effector, suggesting the importance of effectors in modulating host immune responses for successful host colonization.

In this study, we documented the formation of appressoria by *M. anisopliae* during germination on its host. Previously, only *M. robertsii* has been reported to produce “appressorium-like structures” on *Arabidopsis* roots, so called because the germling swollen at the tip of as an appressorium after day one post inoculation of the fungus on the roots (55). We observed the formation of similar structures in *M. anisopliae* during germination on barley roots. Spore germination stages occurred earlier on the barley root surface in *M. anisopliae* compared to growth on PDA, and the appressoria formed 26 hours after inoculation of the roots. Phytohormones, including strigolactones, IAA, and gibberellic acid, have been shown to influence the maintenance of *Metarhizium*– plant associations (56, 57), and have been found to induce conidial germination and hyphal growth in *M. guizhouense* when assayed on media amended with phytohormones (58). Similar results were also reported in mycorrhizal fungi, which exhibit extensive hyphal branching in the proximity of host roots, suggesting a role for plant metabolites in promoting fungal conidial germination and growth during beneficial fungal-plant interactions (59, 60), consistent with our results.

The *F. graminearum* genome harbors more CAZyme-encoding genes than the genome of *M. anisopliae* (Fig.4A and B). This abundance of CAZymes in *F. graminearum*, together with the higher expression levels of these genes during interactions with the host, provide supporting evidence for the enhanced capacity of *F. graminearum* to penetrate the plant host and initiate disease. In contrast, *M. anisopliae* has limited expression of genes for plant cell wall-degrading enzymes during initial colonization, reflecting a less aggressive approach for establishment in the host. In general, fungal colonization is initiated when conidia first adhere to, and then breach the cuticle. Cutinases act as surface-sensing virulence factors and are crucial for establishing infection in above-ground plant parts. Cutinase has been reported as essential in disease development in phytopathogenic fungi including *Botryosphaeria dothidea* (61)*, F. sacchari* (62)*, Magnaporthe oryzae* (63) and *Sclerotinia sclerotiorum* (64) and cutinase genes are upregulated in *F. graminearum* during appressorium formation. We found a lower number of cutinase and polysaccharide lyase genes in the *M. anisopliae* genome than in the *F. graminearum* genome. Additionally, we documented a low germination rate for *M. anisopliae* when inoculated onto barley leaf sheaths (data not shown) when compared to the barley roots. The limited expression of CAZyme genes in *M. anisopliae* allows it to enter the host while minimizing damage, which is consistent with its ability to colonize root cortical cells and intracellular spaces with limited impact on plant health (29).

Our data show that multiple specialized metabolite gene clusters are upregulated during appressorium formation in *F. graminearum* compared to *M. anisopliae*. However, the metabolic pathway analysis of upregulated genes of *M. anisopliae* on the host showed that the tryptophan metabolic pathway was more highly expressed in the appressorium stage. Tryptophan metabolism is linked to the IAA synthesis, and we found that the candidate genes involved in the synthesis of IAA were upregulated in the appressorium stage. *Metarhizium* spp. have been reported to promote root hair growth promotion within 1-2 days of colonization of the host (29). Previous studies showed that IAA influences the growth and architecture of roots, which are important for nutrient and water uptake (65, 66). The ability of *M. anisopliae* to induce root growth in plants through IAA dependent mechanisms (54) demonstrates the intricate and multifaceted relationship that exists between this endophyte and its plant host. This difference highlights how *F. graminearum* uses a wider range of specialized metabolites for aggressive plant colonization, while *M. anisopliae* employs a phytohormone-based strategy to interact with its plant host.

Our results indicate that genes encoding secreted putative effectors are upregulated during the appressorium stage for both species, emphasizing the pivotal role of secreted effector proteins in the early stages of plant colonization, regardless of the lifestyle of the fungus. Secreted effectors are generally described as cysteine-rich proteins with a signal peptide for secretion and have no transmembrane domain, used by plant colonizing fungi to manipulate the host defense responses (67, 68). However, the number of putative effectors expressed was higher in the phytopathogen, *F. graminearum,* compared to *M. anisopliae*, an endophyte and mutualist. We found that putative effector genes, including LysM protein coding genes reported in other plant colonizing fungi (69–71), were upregulated in both *M. anisopliae* and *F. graminearum* during colonization of the host. The LysM domain in the effectors suppresses the chitin-induced plant defense response and has been identified previously in other plant colonizing fungi as modulating host immunity (72). The role of LysM proteins in modulating defense responses in plant hosts has been described in *Botryosphaeria dothidea* (73), *Trichoderma atroviride* (74) and *Verticillium dahliae* (75). The deletion of the *lysm2* gene reduced the ability of beneficial plant root colonizer *Clonostachys rosea* to colonize wheat roots (76). The two LysM proteins, ChELP1 and ChELP2, in the anthracnose fungus *Colletrotrichum higginsianum,* have been shown to be essential for fungal attachment, appressorium function, and suppression of chitin triggered immunity of hosts (77). The differential upregulation of putative LysM type effectors in both *M. anisopliae* and *F. graminearum* during appressorium formation suggests that suppressing chitin-triggered immunity in plant hosts is an important step during plant colonization and is an evolutionarily conserved trait in both pathogenic and mutualistic/endophytic fungal species.

Our analysis suggests that virulence-associated genes are important for both *F. graminearum* and *M. anisopliae* to initiate interactions with plant hosts. For instance, Egh16-like protein coding genes, previously reported in both phytopathogenic and entomopathogenic fungi as virulence factors, were upregulated in both *F. graminearum* and *M. anisopliae* during *in planta* growth. In the rice blast fungus, *M. oryzae*, *Egh16*-like genes have been shown to play a role in host penetration and lesion formation (78). Similarly, in *Metarhizium acridum*, a specialist insect pathogen, the deletion of *Magas1*, which encodes a protein containing an Egh16-like virulence factor domain, impaired fungal penetration during topical infection of locusts (79). Additionally, we observed increased expression of genes encoding necrosis-inducing proteins during the appressorium stage of *F. graminearum*, suggesting the initiation of the necrotrophic phase of *F. graminearum* even at this early stage of infection. The necrosis-inducing protein family has been reported in plant pathogenic bacteria, oomycetes, and pathogenic fungi, triggering cell death and suppressing defense responses on the host (80). For instance, the necrosis-inducing protein (Nep1) of *F. oxysporum* was implicated in triggering cell death and necrotic spots in *Arabidopsis* (81).

We found major differences in the expression profiles of plant cell wall degrading enzymes and specialized metabolites between two fungi that play distinct roles during infection of their hosts. *F. graminearum* expresses multiple CAZyme genes and specialized metabolite gene clusters that have been linked to its pathogenic lifestyle, whereas *M. anisopliae* expresses genes associated with phytohormones. Each approach plays a key role in adaptation to a specific lifestyle. Future research should provide an understanding of the functional roles of the CAZymes, effectors, and specialized metabolites identified in this comparative analysis. Additionally, expanding this research to include proteomic and metabolomic approaches will provide a more comprehensive understanding of the underlying biochemical processes of spore germination and adaptive mechanisms employed by these fungi during host interactions. Elucidation of these processes will be critical to developing targeted strategies for managing fungal diseases and for optimizing applications of beneficial fungi.

## ACKNOWLEDGEMENTS

This work was supported by the US Department of Agriculture National Institute of Food and Agriculture under Award No. 2020-67013-31185 to FT. We acknowledge the support of Michigan State University AgBioResearch. Many thanks to Dr. Carolyn Young, who was involved in conceptualization of the project and procuring the funding, and then moved on to chair a department.

## SUPPLEMENTARY MATERIAL

### Supplementary Figures

**Fig. S1** Volcano plots of differentially expressed genes (DEGs) from the transcriptome data of *F. graminearum*. The number of up- or down-regulated DEGs of *F. graminearum* in Stages 2, 3 and 4 vs Stage 1 on (**A-C**) barley and (**D-F**) PDA are shown.

**Fig. S2** Volcano plots of DEGs from the transcriptome data of *M. anisopliae*. The number of up- or down-regulated DEGs of *M. anisopliae* in Stages 2, 3 and 4 vs Stage 1 on (**A-C**) barley and (**D-F**) PDA are shown.

**Fig. S3** Top 100 genes of (**A**) *F. graminearum* and (**B**) *M. anisopliae* showed highly variable expression levels during different germination stages on barley and PDA. Genes were grouped into five clusters based on the expression pattern. The heatmap color range represents high to low expression levels (rlog-transformed normalized counts) where red represents higher expression and blue represents lower expression.

**Fig. S4** Gene ontology classification of DEGs of *F. graminearum* in different germination stages on (**A-C**) barley and (**D-F**) PDA. The numbers on the bar represent the quantity of upregulated genes (LFC ≥ 5 and adjusted p-adj ≤ 0.01) enriched for each GO term.

**Fig. S5** Gene ontology classification of DEGs of *M. anisopliae* in different germination stages on (**A-C**) barley and (**D-F**) PDA. The numbers on the bar represent the quantity of upregulated genes (LFC ≥ 5 and adjusted p-adj ≤ 0.01) enriched for each GO term.

**Fig. S6** The shared metabolic pathways between *F. graminearum* and *M. anisopliae* for each germination stage on both nutritional conditions (Stages 2, 3 and 4). (**A**) Stage 2 vs 1. (**B**) Stage 3 vs 1. (**C**) Stage 4 vs 1.

**Fig. S7** Expression of trichothecene biosynthetic genes during different germination stages in *F. graminearum* on barley and PDA. Heatmap colour range represents high to low expression levels where the red represents higher expression and blue represents lower expression.

### Supplementary Tables

**Table S1.** Differential expression analysis of *F. graminearum* in each germination stage (Stages 2, 3 and 4) on barley and PDA.

**Table S2.** Differential expression analysis of *M. anisopliae* in each germination stage (Stages 2, 3 and 4) on barley and PDA.

**Table S3.** GO enrichment analysis of differentially expressed upregulated genes (LFC ≥ 5 and p-adj ≤0.01) of *F. graminearum* in each germination stage (Stages 2, 3 and 4) on barley and PDA.

**Table S4.** GO enrichment analysis of differentially expressed upregulated genes (LFC ≥ 5 and p-adj ≤0.01) of *M. anisopliae* in each germination stage (Stages 2, 3 and 4) on barley and PDA.

**Table S5.** Metabolic pathway analysis of differentially expressed upregulated genes (LFC ≥ 5 and p-adj ≤0.01) of *F. graminearum* in each germination stage (Stages 2, 3 and 4) on barley and PDA.

**Table S6.** Metabolic pathway analysis of differentially expressed upregulated genes (LFC ≥ 5 and p-adj ≤0.01) of *M. anisopliae* in each germination stage (Stages 2, 3 and 4) on barley and PDA.

**Table S7.** The common metabolic pathways for the appressoria stage in *F. graminearum* and *M. anisopliae*.

**Table S8.** CAZymes identified in *F. graminearum.*

**Table S9.** CAZymes identified in *M. anisopliae.*

**Table S10.** Specialized metabolite gene clusters identified in *F. graminearum.*

**Table S11.** Specialized metabolite gene clusters identified in *M. anisopliae.*

**Table S12.** Expression analysis of putative genes involved in IAA biosynthesis in *M. anisopliae*.

**Table S13.** Putative effectors identified in *F. graminearum*.

**Stable 14.** Putative effectors identified in *M. anisopliae*.

## REFERENCES

1. Rgen Wendland J. 2001. Comparison of morphogenetic networks of filamentous fungi and yeast. Fungal Genet Biol 34:63–82.

2. Seong KY, Zhao X, Xu JR, Güldener U, Kistler HC. 2008. Conidial germination in the filamentous fungus *Fusarium graminearum*. Fungal Genet Biol 45:389–399.

3. van Leeuwen MR, Krijgsheld P, Bleichrodt R, Menke H, Stam H, Stark J, Wösten HAB, Dijksterhuis J. 2013. Germination of conidia of *Aspergillus niger* is accompanied by major changes in RNA profiles. Stud Mycol 74:59–70.

4. Ryder LS, Talbot NJ, Paszkowski U, Scott B. 2015. Regulation of appressorium development in pathogenic fungi. Curr Opin Plant Biol 26:8–13.

5. Osherov N, May GS. 2001. The molecular mechanisms of conidial germination. FEMS Microbiol Lett 199:153–160.

6. Li X, Li B hua, Lian S, Dong X li, Wang C xia, Liang W xing. 2019. Effects of temperature, moisture and nutrition on conidial germination, survival, colonization and sporulation of *Trichothecium roseum*. Eur J Plant Pathol 153:557–570.

7. Ijadpanahsaravi M, Punt M, Wösten HAB, Teertstra WR. 2021. Minimal nutrient requirements for induction of germination of *Aspergillus niger* conidia. Fungal Biol 125:231–238.

8. Ji T, Altieri V, Salotti I, Rossi V. 2023. Effects of temperature and moisture duration on spore germination of four fungi that cause grapevine trunk diseases. Plant Dis 107:1005– 1008.

9. Mendgen K, Hahn M, Deising H. 1996. Morphogenesis and mechanisms of penetration by plant pathogenic fungi. Annu Rev Phytopathol 12:367–86.

10. Pedrini N, Ortiz-Urquiza A, Huarte-Bonnet C, Zhang S, Keyhani NO. 2013. Targeting of insect epicuticular lipids by the entomopathogenic fungus *Beauveria bassiana*: Hydrocarbon oxidation within the context of a host-pathogen interaction. Front Microbiol 4:1–18.

11. Chethana KWT, Jayawardena RS, Chen YJ, Konta S, Tibpromma S, Abeywickrama PD, Gomdola D, Balasuriya A, Xu J, Lumyong S, Hyde KD. 2021. Diversity and function of appressoria. Pathogens 10:3–6.

12. Sikhakolli UR, López-Giráldez F, Li N, Common R, Townsend JP, Trail F. 2012. Transcriptome analyses during fruiting body formation in *Fusarium graminearum* and *Fusarium verticillioides* reflect species life history and ecology. Fungal Genet Biol 49:663–673.

13. Roux J, Rosikiewicz M, Robinson-Rechavi M. 2015. What to compare and how: Comparative transcriptomics for Evo-Devo. J Exp Zool Part B Mol Dev Evol 324:372– 382.

14. Trail F, Wang Z, Stefanko K, Cubba C, Townsend JP. 2017. The ancestral levels of transcription and the evolution of sexual phenotypes in filamentous fungi. PLoS Genet 13:e1006867.

15. Chan ME, Bhamidipati PS, Goldsby HJ, Hintze A, Hofmann HA, Young RL. 2021. Comparative transcriptomics reveals distinct patterns of gene expression conservation through vertebrate embryogenesis. Genome Biol Evol 13.

16. Miguel-Rojas C, Cavinder B, Townsend JP, Trail F. 2023. Comparative transcriptomics of *Fusarium graminearum* and *Magnaporthe oryzae* spore germination leading up to infection. MBio 14.

17. Zeilinger S, Gupta VK, Dahms TES, Silva RN, Singh HB, Upadhyay RS, Gomes EV, Tsui CKM, Chandra Nayak S. 2016. Friends or foes? Emerging insights from fungal interactions with plants. FEMS Microbiol Rev 40:182–207.

18. Pontes JGDM, Fernandes LS, Dos Santos R Vander, Tasic L, Fill TP. 2020. Virulence factors in the phytopathogen-host interactions: An overview. J Agric Food Chem 68:7555–7570.

19. König A, Müller R, Mogavero S, Hube B. 2021. Fungal factors involved in host immune evasion, modulation and exploitation during infection. Cell Microbiol 23:e13272.

20. Pradhan A, Ghosh S, Sahoo D, Jha G. 2021. Fungal effectors, the double edge sword of phytopathogens. Curr Genet 67:27–40.

21. Westrick NM, Smith DL, Kabbage M. 2021. Disarming the host: Detoxification of plant defense compounds during fungal necrotrophy. Front Plant Sci 12:651716.

22. Gutiérrez-Sánchez A, Plasencia J, Monribot-Villanueva JL, Rodríguez-Haas B, Ruíz-May E, Guerrero-Analco JA, Sánchez-Rangel D. 2023. Virulence factors of the genus *Fusarium* with targets in plants. Microbiol Res 277:127506.

23. Plett JM, Martin F. 2015. Reconsidering mutualistic plant–fungal interactions through the lens of effector biology. Curr Opin Plant Biol 26:45–50.

24. Patel P, Kumar S, Modi A, Kumar A. 2021. Deciphering fungal endophytes combating abiotic stresses in crop plants (cereals and vegetables). Microb Manag Plant Stress Curr Trends, Appl Challenges 131–147.

25. Goswami RS, Kistler HC. 2004. Heading for disaster: *Fusarium graminearum* on cereal crops. Mol Plant Pathol 5:515–525.

26. Osborne LE, Stein JM. 2007. Epidemiology of Fusarium head blight on small-grain cereals. Int J Food Microbiol 119:103–108.

27. Trail F. 2009. For blighted waves of grain: *Fusarium graminearum* in the postgenomics era. Plant Physiol 149:103–110.

28. Hu G, St Leger RJ. 2002. Field studies using a recombinant mycoinsecticide (*Metarhizium anisopliae*) reveal that it is rhizosphere competent. Appl Environ Microbiol 68:6383– 6387.

29. Sasan RK, Bidochka MJ. 2012. The insect-pathogenic fungus *Metarhizium robertsii* (Clavicipitaceae) is also an endophyte that stimulates plant root development. Am J Bot 99:101–107.

30. Zimmermann G. 1993. The entomopathogenic fungus *Metarhizium anisopliae* and its potential as a biocontrol agent. Pestic Sci 37:375–379.

31. Roberts DW, Leger RJS. 2004. *Metarhizium* spp., cosmopolitan insect-pathogenic fungi: Mycological aspects. Adv Appl Microbiol 54:1–70.

32. Behie SW, Bidochka MJ. 2014. Ubiquity of insect-derived nitrogen transfer to plants by endophytic insect-pathogenic fungi: an additional branch of the soil nitrogen cycle. Appl Environ Microbiol 80:1553–60.

33. Behie SW, Moreira CC, Sementchoukova I, Barelli L, Zelisko PM, Bidochka MJ. 2017. Carbon translocation from a plant to an insect-pathogenic endophytic fungus. Nat Commun 8:14245.

34. Bolger AM, Lohse M, Usadel B. 2014. Trimmomatic: A flexible trimmer for Illumina sequence data. Bioinformatics 30:2114–2120.

35. King R, Urban M, Hammond-Kosack MCU, Hassani-Pak K, Hammond-Kosack KE. 2015. The completed genome sequence of the pathogenic ascomycete fungus *Fusarium graminearum*. BMC Genomics 16:1–21.

36. Hu X, Xiao G, Zheng P, Shang Y, Su Y, Zhang X, Liu X, Zhan S, St. Leger RJ, Wang C. 2014. Trajectory and genomic determinants of fungal-pathogen speciation and host adaptation. Proc Natl Acad Sci 111:16796–16801.

37. Kim D, Langmead B, Salzberg SL. 2015. HISAT: A fast spliced aligner with low memory requirements. Nat Methods 12:357–360.

38. Li H, Handsaker B, Wysoker A, Fennell T, Ruan J, Homer N, Marth G, Abecasis G, Durbin R. 2009. The sequence alignment/map format and SAMtools. Bioinformatics 25:2078–2079.

39. Anders S, Pyl PT, Huber W. 2015. HTSeq-A Python framework to work with high-throughput sequencing data. Bioinformatics 31:166–169.

40. Zhang H, Yohe T, Huang L, Entwistle S, Wu P, Yang Z, Busk PK, Xu Y, Yin Y. 2018. dbCAN2: a meta server for automated carbohydrate-active enzyme annotation. Nucleic Acids Res 46:W95–W101.

41. Blin K, Shaw S, Kloosterman AM, Charlop-Powers Z, Van Wezel GP, Medema MH, Weber T. 2021. antiSMASH 6.0: improving cluster detection and comparison capabilities. Nucleic Acids Res 49:W29–W35.

42. Gaffoor I, Brown DW, Plattner R, Proctor RH, Qi W, Trail F. 2005. Functional analysis of the polyketide synthase genes in the filamentous fungus *Gibberella zeae* (Anamorph *Fusarium graminearum*). Eukaryot Cell 4:1926–1933.

43. Sieber CMK, Lee W, Wong P, Mü Nsterkö Tter M, Mewes H-W. 2014. The *Fusarium graminearum* genome reveals more secondary metabolite gene clusters and hints of horizontal gene transfer. PLoS One 9:110311.

44. Mentges M, Glasenapp A, Boenisch M, Malz S, Henrissat B, Frandsen RJN, Güldener U, Münsterkötter M, Bormann J, Lebrun MH, Schäfer W, Martinez-Rocha AL. 2020. Infection cushions of *Fusarium graminearum* are fungal arsenals for wheat infection. Mol Plant Pathol 21:1070–1087.

45. Hicks C, Witte TE, Sproule A, Hermans A, Shields SW, Colquhoun R, Blackman C, Boddy CN, Subramaniam R, Overy DP. 2023. CRISPR-Cas9 gene editing and secondary metabolite screening confirm *Fusarium graminearum* C16 biosynthetic gene cluster products as decalin-containing diterpenoid pyrones. J Fungi 9:695.

46. Sperschneider J, Dodds PN, Gardiner DM, Singh KB, Taylor JM. 2018. Improved prediction of fungal effector proteins from secretomes with EffectorP 2.0. Mol Plant Pathol 19:2094–2110.

47. Harris MA, Clark J, Ireland A, Lomax J, Ashburner M, Foulger R, Eilbeck K, Lewis S, Marshall B, Mungall C, Richter J, Rubin GM, Blake JA, Bult C, Dolan M, Drabkin H, Eppig JT, Hill DP, Ni L, Ringwald M, Balakrishnan R, Cherry JM, Christie KR, Costanzo MC, Dwight SS, Engel S, Fisk DG, Hirschman JE, Hong EL, Nash RS, Sethuraman A, Theesfeld CL, Botstein D, Dolinski K, Feierbach B, Berardini T, Mundodi S, Rhee SY, Apweiler R, Barrell D, Camon E, Dimmer E, Lee V, Chisholm R, Gaudet P, Kibbe W, Kishore R, Schwarz EM, Sternberg P, Gwinn M, Hannick L, Wortman J, Berriman M, Wood V, de la Cruz N, Tonellato P, Jaiswal P, Seigfried T, White R. 2004. The Gene Ontology (GO) database and informatics resource. Nucleic Acids Res 32:D258–D261.

48. Caspi R, Billington R, Ferrer L, Foerster H, Fulcher CA, Keseler IM, Kothari A, Krummenacker M, Latendresse M, Mueller LA, Ong Q, Paley S, Subhraveti P, Weaver DS, Karp PD. 2015. The MetaCyc database of metabolic pathways and enzymes and the BioCyc collection of pathway/genome databases. Nucleic Acids Res 44:471–480.

49. Stajich JE, Harris T, Brunk BP, Brestelli J, Fischer S, Harb OS, Kissinger JC, Li W, Nayak V, Pinney DF, Stoeckert CJ, Roos DS. 2012. FungiDB: an integrated functional genomics database for fungi. Nucleic Acids Res 40:D675–D681.

50. Fan X, Zhang P, Batool W, Liu C, Hu Y, Wei Y, He Z, Zhang SH. 2023. Contribution of the tyrosinase (MoTyr) to melanin synthesis, conidiogenesis, appressorium development, and pathogenicity in *Magnaporthe oryzae*. J Fungi 9:311.

51. Zhao Y. 2012. Auxin biosynthesis: A simple two-step pathway converts tryptophan to indole-3-acetic acid in plants. Mol Plant 5:334–338.

52. Wang B, Chu J, Yu T, Xu Q, Sun X, Yuan J, Xiong G, Wang G, Wang Y, Li J. 2015. Tryptophan-independent auxin biosynthesis contributes to early embryogenesis in *Arabidopsis*. Proc Natl Acad Sci U S A 112:4821–4826.

53. Solanki M, Shukla LI. 2023. Recent advances in auxin biosynthesis and homeostasis. 3 Biotech 2023 139 13:1–22.

54. Liao X, Lovett B, Fang W, St Leger RJ. 2017. *Metarhizium robertsii* produces indole-3-acetic acid, which promotes root growth in *Arabidopsis* and enhances virulence to insects. Microbiology 163:980–991.

55. Dai J, Tang X, Wu C, Liu S, Mi W, Fang W. 2024. Utilization of plant-derived sugars and lipids are coupled during colonization of rhizoplane and rhizosphere by the fungus *Metarhizium robertsii*. Fungal Genet Biol 172:103886.

56. Hu S, Bidochka MJ. 2021. Abscisic acid implicated in differential plant responses of *Phaseolus vulgaris* during endophytic colonization by *Metarhizium* and pathogenic colonization by *Fusarium*. Sci Reports 2021 111 11:1–12.

57. Ahmad I, del Mar Jiménez-Gasco M, Luthe DS, Barbercheck ME. 2022. Endophytic *Metarhizium robertsii* suppresses the phytopathogen, *Cochliobolus heterostrophus* and modulates maize defenses. PLoS One 17:e0272944.

58. Piña-Torres IH, Dávila-Berumen F, González-Hernández GA, Torres-Guzmán JC, Padilla-Guerrero IE. 2023. Hyphal growth and conidia germination are induced by phytohormones in the root colonizing and plant growth promoting fungus *Metarhizium guizhouense*. J Fungi 9:945.

59. Barker SJ, Tagu D. 2000. The roles of auxins and cytokinins in mycorrhizal symbioses. J Plant Growth Regul 19:144–154.

60. Akiyama K, Ogasawara S, Ito S, Hayashi H. 2010. Structural requirements of strigolactones for hyphal branching in AM fungi. Plant Cell Physiol 51:1104–1117.

61. Dong BZ, Zhu XQ, Fan J, Guo LY. 2021. The cutinase *Bdo_10846* play an important role in the virulence of *Botryosphaeria dothidea* and in inducing the wart symptom on apple plant. Int J Mol Sci 22:1910.

62. Liang H, Li F, Huang Y, Yu Q, Huang Z, Zeng Q, Chen B, Meng J. 2024. FsCGBP, a cutinase G-Box binding protein, regulates the growth, development, and virulence of *Fusarium sacchari*, the pathogen of sugarcane Pokkah boeng disease. J Fungi 10:246.

63. Skamnioti P, Gurr SJ. 2007. *Magnaporthe grisea* cutinase2 mediates appressorium differentiation and host penetration and is required for full virulence. Plant Cell 19:2674– 2689.

64. Gong Y, Fu Y, Xie J, Li B, Chen T, Lin Y, Chen W, Jiang D, Cheng J. 2022. *Sclerotinia sclerotiorum* SsCut1 modulates virulence and cutinase activity. J Fungi 8:526.

65. Chen Q, Dai X, De-Paoli H, Cheng Y, Takebayashi Y, Kasahara H, Kamiya Y, Zhao Y. 2014. Auxin overproduction in shoots cannot rescue auxin deficiencies in *Arabidopsis* roots. Plant Cell Physiol 55:1072–1079.

66. Liu R, Yang L, Zou Y, Wu Q. 2023. Root-associated endophytic fungi modulate endogenous auxin and cytokinin levels to improve plant biomass and root morphology of trifoliate orange. Hortic Plant J 9:463–472.

67. Jaswal R, Kiran K, Rajarammohan S, Dubey H, Singh PK, Sharma Y, Deshmukh R, Sonah H, Gupta N, Sharma TR. 2020. Effector biology of biotrophic plant fungal pathogens: Current advances and future prospects. Microbiol Res 241:126567.

68. Shao D, Smith DL, Kabbage M, Roth MG. 2021. Effectors of plant necrotrophic fungi. Front Plant Sci 12:687713.

69. Kombrink A, Thomma BPHJ. 2013. LysM effectors: Secreted proteins supporting fungal life. PLOS Pathog 9:e1003769.

70. Lee WS, Rudd JJ, Hammond-Kosack KE, Kanyuka K. 2014. *Mycosphaerella graminicola* lysm effector-mediated stealth pathogenesis subverts recognition through both cerk1 and cebip homologues in wheat. Mol Plant-Microbe Interact 27:236–243.

71. Hu SP, Li JJ, Dhar N, Li JP, Chen JY, Jian W, Dai XF, Yang XY. 2021. Lysin motif (LysM) proteins: Interlinking manipulation of plant immunity and fungi. Int J Mol Sci 22:3114.

72. Dölfors F, Holmquist L, Dixelius C, Tzelepis G. 2019. A LysM effector protein from the basidiomycete *Rhizoctonia solani* contributes to virulence through suppression of chitin-triggered immunity. Mol Genet Genomics 294:1211–1218.

73. Zhang H, Wen SH, Li PH, Lu LY, Yang X, Zhang CJ, Guo LY, Wang D, Zhu XQ. 2023. LysM protein BdLM1 of *Botryosphaeria dothidea* plays an important role in full virulence and inhibits plant immunity by binding chitin and protecting hyphae from hydrolysis. Front Plant Sci 14:1320980.

74. Romero-Contreras YJ, Ramírez-Valdespino CA, Guzmán-Guzmán P, Macías-Segoviano JI, Villagómez-Castro JC, Olmedo-Monfil V. 2019. Tal6 from *Trichoderma atroviride* is a LysM effector involved in mycoparasitism and plant association. Front Microbiol 10:479815.

75. Kombrink A, Rovenich H, Shi-Kunne X, Rojas-Padilla E, van den Berg GCM, Domazakis E, de Jonge R, Valkenburg DJ, Sánchez-Vallet A, Seidl MF, Thomma BPHJ. 2017. *Verticillium dahliae* LysM effectors differentially contribute to virulence on plant hosts. Mol Plant Pathol 18:596–608.

76. Dubey M, Vélëz H, Broberg M, Jensen DF, Karlsson M. 2020. LysM proteins regulate fungal development and contribute to hyphal protection and biocontrol traits in *Clonostachys rosea*. Front Microbiol 11.

77. Takahara H, Hacquard S, Kombrink A, Hughes HB, Halder V, Robin GP, Hiruma K, Neumann U, Shinya T, Kombrink E, Shibuya N, Thomma BPHJ, O’Connell RJ. 2016. *Colletotrichum higginsianum* extracellular LysM proteins play dual roles in appressorial function and suppression of chitin-triggered plant immunity. New Phytol 211:1323–1337.

78. Xu JR, Xue C. 2002. Time for a blast: genomics of *Magnaporthe grisea*. Mol Plant Pathol 3:173–176.

79. Cao Y, Zhu X, Jiao R, Xia Y. 2012. The Magas1 gene is involved in pathogenesis by affecting penetration in *Metarhizium acridum*. J Microbiol Biotechnol 22:889–893.

80. Pemberton CL, Salmond GPC. 2004. The Nep1-like proteins—a growing family of microbial elicitors of plant necrosis. Mol Plant Pathol 5:353–359.

81. Bae H, Kim MS, Sicher RC, Bae HJ, Bailey BA. 2006. Necrosis- and ethylene-inducing peptide from *Fusarium oxysporum* induces a complex cascade of transcripts associated with signal transduction and cell death in Arabidopsis. Plant Physiol 141:1056–1067.

